# Gender and geographical disparity in editorial boards of journals in psychology and neuroscience

**DOI:** 10.1101/2021.02.15.431321

**Authors:** Eleanor R. Palser, Maia Lazerwitz, Aikaterini Fotopoulou

## Abstract

While certain metrics of diversity have seen great improvement in recent years in academic psychology and neuroscience, unequal representation remains for many positions of power. Here, we reviewed publicly available information in order to infer the proportion of editors by gender and their country of affiliation in the top 50 journals worldwide in each of the two fields. The sample included a total of 2,864 editors for psychology journals and 3,093 editors for neuroscience journals. There was a statistically significant difference in the proportion of male and female editors in both fields, both across editorial roles, and within various role categories, including editor-in-chief and their deputies at neuroscience journals, associate and section editors in both fields, and editorial and advisory board members in both fields. The only category in which there was not a significant imbalance of male and female scholars was the editors-in-chief of psychology journals and their deputies. Geographically, USA-based academics significantly outnumbered those from other countries as editors in both psychology and neuroscience. Results also indicated that over three quarters of psychology journals (76%) were comprised of more than 50% male editors, while only 20% had a similar proportion of female editors. In neuroscience, 88% of journals were comprised of more than 50% male editors, while only 10% of journals included a similar, proportional majority of female editors. Findings suggest that editorial positions in academic journals, possibly one of the most powerful decision-making roles in academic psychology and neuroscience, are not balanced in gender or geographical representation.

## Introduction

The landscape of psychology and neuroscience has changed dramatically over the past century with respect to gender, race and nationality, however, important gaps remain in career advancement, particularly in the later stages of career attainment (Gruber et al., 2020; Roberts, Bareket-Shavit, Dollins, Goldie, & Mortenson, 2020; Ryu, 2010). While gender gaps in tenure-track hiring decisions and promotion rates appear to have narrowed, evidence suggests female academics remain under-represented in senior career phases. For example, in data from the USA and Canada, females outnumber males by approximately three to one in psychology graduate programs and comprise approximately half of neuroscience graduate programs, and have done so for more than a decade (Fowler et al., 2018; Stricker, 2003). In the USA, female early career scholars are less likely than males to apply for tenure-track positions (Webber & Gonzales Canché, 2018), however, they are equally as likely, and perhaps even more likely, to be hired when they do (Ginther & Kahn, 2014; National Research Council, 2010; Williams & Ceci, 2015). Female scholars, however, are under-represented among the ranks of full professors (Ginther & Kahn, 2014) and earn, on average, 88% of what their male peers in these roles do (National Science Foundation [NSF], 2018).

Positions of power and indicators of eminence also persist as areas of inequality (Eagly & Miller, 2016). Women are under-represented amongst the ranks of public intellectuals, comprising, for example, only a quarter of authors listed in The New York Times’ *Grey Matter section* (Gruber et al., 2020). Previous reports in the last decade have found that the editorial boards of leading scientific journals in medicine, and the subfields of psychiatry and neurology, feature significantly more men than women (Amrein, Langmann, Fahrleitner-Pammer, Pieber, & Zollner-Schwetz, 2011; Hafeez et al., 2019; Mariotto, Beatrice, Carta, Silvia, Bozzetti, & Mantovani et al., 2020). To our knowledge, the fields of neuroscience and psychology have not been subjected to such an analysis (although see Gruber et al., 2020, for an analysis of American Psychological Association [APA] and Association for Psychological Science [APS] journals).

Journal editors exert considerable power over what is published, and by extension, the direction of an academic discipline and the career advancement of authors. It is important then, to minimize any biases extrinsic to the merit of the work that affect publication decisions. One way to achieve this to ensure a diverse pool of editors, such that biases are diluted, and their influence reduced. Internationalization has been cited as an important goal in achieving diversity and innovation (Weick, 1995). Worryingly, previous research suggests that the geographical representation of journal article authors is strongly associated with that of the editorial boards (Baruch, 2001; Granadino, Garcia-Carpintero, & Plaza, 2006). In an analysis of the editorial boards of the top 20 journals in 15 scientific disciplines, of which neuroscience was one, a significant logarithmic relationship was observed between the nationalities of editorial board members, and the number of publications originating from those countries (Garcia-Carpintero, Granadino & Plaza, 2010). While the directionality of these findings is difficult to ascertain, they highlight the potential for bias and hegemony in academic publishing and suggest that in addition to gender, the geographical representation of editors is another important factor to consider when quantifying disparity in academic publishing.

Here, we consider the top 50 journals in psychology and neuroscience, as ranked by an independent source, Science Citation Index Expanded (SCIE) list in *Clarivate Analytics’ Journal Citation Reports* (JCR), in terms of the gender and geographical affiliation of their editors. We consider different categories of editor, in line with similar work in psychiatry (Hafeez et al., 2019), in order to understand any differences based on relative decision-making power and role. The goal is to provide quantitative data on the current status of editorial positions in these related fields, that can be used to monitor progress over time and act as a starting point for a deeper, quantitative and qualitative understanding of the reasons for any uneven representation, and remedial action.

## Methods

The top 50 journals for the year 2019 in the fields of “psychology” and “neuroscience” were selected from the Science Citation Index Expanded (SCIE) list in *Clarivate Analytics’ Journal Citation Reports* (JCR). Selection was limited to journals published in the English language. This resulted in two databases of 50 editorial boards each (see Supplementary data 1 and 2 for psychology and neuroscience databases, respectively). Note that the list generated for psychology contains the top 50 generalist journals in psychology, and as such does not contain high-impact journals within sub-disciplines of psychology. There was no overlap in the top 50 journals for the two fields.

### Data collection

Journal webpages were audited for editorial board members. Manual data inspection and entry was conducted in October and November 2020, with a final check of the databases completed in the first week of December, 2020. Any changes to editorial boards made after this time were not included in the databases. Each journal’s country of publication was downloaded from the JCR. The majority of the journals were published in the United States and United Kingdom. For psychology, 58% of journals were published in the US, and 24% were published in the UK. For neuroscience, 46% of journals were published in the UK, and 40% were published in the US. See Supplementary data 1 and 2 for the country of publication of all journals included.

Certain editorial roles were not included in the databases as these were deemed not to be decision-making roles (e.g., managing editors, student advisors, social media editors). See Supplementary Table 1 for a full list of excluded categories. One journal in the field of psychology, *Journals of Gerontology Series B-Psychological Sciences and Social Sciences*, listed separate editorial boards for the psychological and social sciences. Due to our focus on psychology and neuroscience, we only included the editors for psychological sciences in the database.

Names, role, country of affiliation and gender were manually tabulated based primarily on information used in journals’ public biographical sections for each editor. Pronouns were used as the most reliable indicator of gender, including non-binary gender identities, but when this was not available, we had to make (arguably imprecise inferences based on names and/or images. We acknowledge this is a major limitation of our work as our results are based on inferences based on language and other external characteristics rather than self-descriptions. When gender and/or country of affiliation were not available on journal webpages, an internet search was conducted for these details, browsing institutional webpages, Google Scholar, etc. When gender or country of affiliation could not be discerned, they were marked as “not available” (NA; n = 1 in psychology and n = 8 in neuroscience for gender).

Most journals had fewer than 150 editors (*M* = 156.97; *SD* = 972.61) with the exception of *Frontiers in Psychology*, which was a clear outlier (3 standard deviations above the mean number of editors), with a total n of 9,780 editors. So, in this case, we selected only the Field Chief Editor and Specialty Chief Editors (n = 40).

In line with similar research on the editorial boards of psychiatry journals (Hafeez et al., 2019) editors were categorized according to their role. Initially four categories, exactly mirroring Hafeez et al. (2019), were used: 1) Editor-in-chief and deputies, 2) associate and section editors, 3) editorial board members and 4) advisory board members. The same title was used by different journals to denote varying levels of seniority, and so features in multiple categories. In these instances, decisions were made on an individual basis according on the organization of each journal. See Supplementary Table 1 for the roles assigned to each category. Ultimately, due to the infrequency of journals possessing both an editorial board and an advisory board, categories 3 and 4 were collapsed. As such, the final categorization was as follows: 1) Editor-in-chief and deputies, 2) associate and section editors, and 3) advisory and editorial board members.

Countries of affiliation were classified into six continents according to geographical location: North America (encompassing the United States, Canada and Mexico), Europe (encompassing the United Kingdom, continental Europe, and Russia), Oceania (encompassing Australia and New Zealand), Latin America (encompassing central and south America), Asia (including Turkey and the Middle East) and Africa.

### Statistical analyses

Analyses were conducted separately for psychology and neuroscience journals. One sample Chi-squared tests were used to determine if there were significant differences in the proportion of male and female editors overall in each field and in each of the three editorial categories, irrespective of the proportion of male and female editors at each individual journal. Any editors where the gender could not be reliably discerned were excluded, thus one editor was excluded from gender analyses in the field of psychology, leaving a final sample of 2,863 editors and 8 editors were excluded from gender analyses in neuroscience, leaving a final sample of 3,085 editors. All were from Category 3, advisory and editorial board members. Because of the large number of editors in this category, we are confident that the results would not change if the gender of these editors was known.

In a second analysis, where we considered differences in gender balance between the journals, we calculated what proportion of journals had distributions of male and female editors in ten-point percentage increments from 0 – 100%.

One-sample Chi-squared tests were also used to determine if there were significant differences in the proportion of editors affiliated with each continent, and country. We then quantified how many journals were comprised primarily of editors based in the USA.

Finally, we combined geographical and gender-based data, and quantified the proportion of male and female editors deriving from the major contributing countries. Following inspection of the geographical distribution of our data, we defined a major contributing country as any that contributed ≥4% of total number of editors in a field.

## Results

### Psychology

#### Gender

The sample included a total of 2,864 editors in the field of psychology. We found that, overall, there were significantly more male (n = 1,706) than female (n = 1,157) editors (see **Table 1**). This was driven mainly by the larger categories, namely Category 2, associate and section editors, and Category 3, advisory and editorial board members. There were no significant differences in gender representation amongst the smallest and most senior category, Editors-in-chief and their deputies, although there were more men (n = 49) than women (n = 37) in that category too. This analysis did not take into account variability in the proportion of male and female editors at the individual journals.

**Table 1:** Overall proportion of editors who are male and female in the top 50 journals in the field of psychology, and in each of the three sub-categories: 1) Editors-in-chief and their deputies, 2) associate and section editors, and 3) advisory and editorial boards.

| PSYCHOLOGY | Female | Male | Statistic |
| --- | --- | --- | --- |
| Overall ($n = 2,863$, 1 NA excluded) | 40% | 60% | $\chi^2(1) = 104.9, p < .001$ |
| Editors-in-chief and deputies ($n = 86$) | 43% | 57% | $\chi^2(1) = 1.4, p = .237$ |
| Associate and section editors ($n = 308$) | 44% | 56% | $\chi^2(1) = 5.9, p = .015$ |
| Advisory and editorial boards ($n = 2,368$, 1 NA excluded) | 40% | 60% | $\chi^2(1) = 99.5, p < .001$ |

To account for this, we calculated what percentage of journals had proportions of male and female editors in 10-point percentage increments. Over three quarters of psychology journals (76%) were comprised of more than 50% male editors, while only 20% had a similar proportion of female editors (see **Figure 1A**). Over half of the journals (54%) were comprised of more than 60% male editors, whereas only 8% showed a similar proportion of female editors. Nearly a quarter of journals (22%) were comprised of more than 70% male editors, but only 2% showed a similar proportion of female editors. A similar proportion of journals (2%) were comprised of either more than 80% male or female editors, and more than 90% male (2%) or female (0%) editors. See Supplementary Figure 1 for the binned data by gender at the 50 journals.

**Figure 1:**
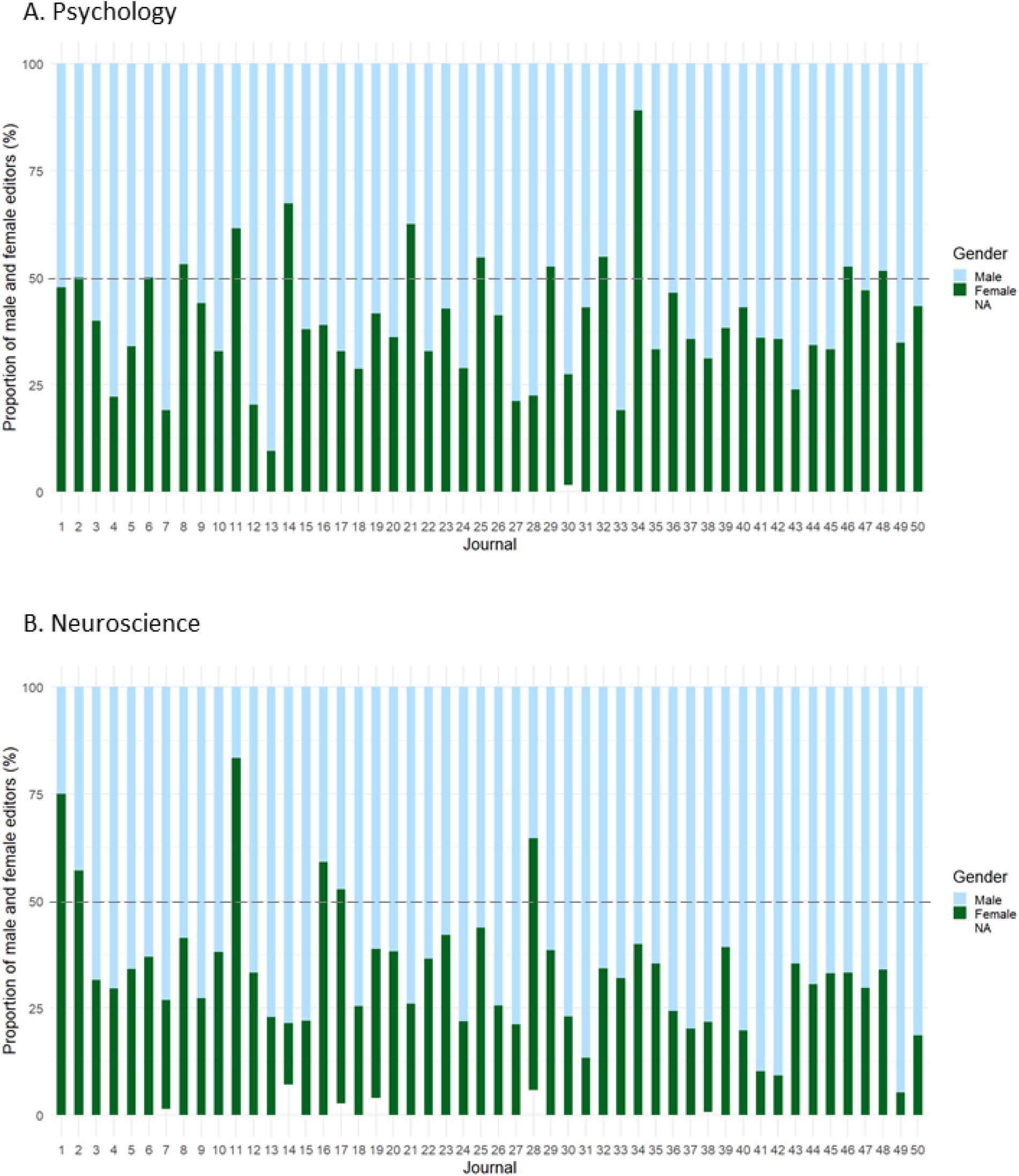
Proportion of male and female editors at the top 50 journals in the fields of psychology (A) and neuroscience (B). Dashed grey horizontal line signifies equal number of male and female editors.

#### Geography

Overall, the editors of the top 50 journals in psychology were primarily based in North America (65%), then, in decreasing order: Europe (26%), Asia (4%) and Oceania (4%), and finally Africa (0.5%) and Latin America (0.5%). This distribution was significantly skewed towards North America [*X*^2^(5) = 3973.2, *p* < .001]. In terms of the country of affiliation, more than half of editors were based in the United States (61%), followed by the United Kingdom (7%), Canada (5%) and Spain (5%). See Supplementary Table 2 for the proportion of editors from all countries that featured. **Figure 2A** shows the proportion of editors from each country that contributed more than 1% to the total number of editors in the field’s top 50 journals.

**Figure 2:**
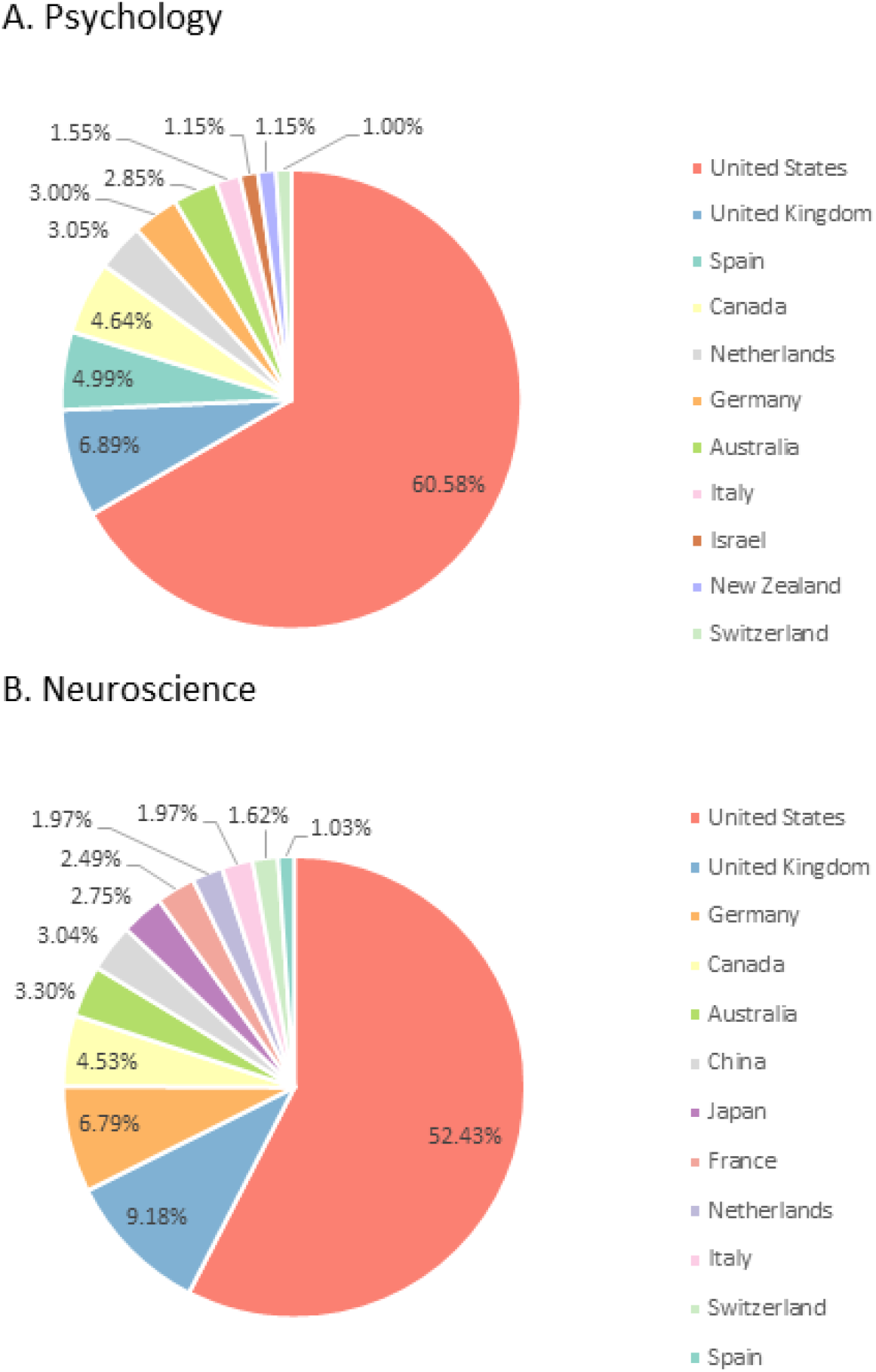
Proportion of editors at the top 50 journals in the fields of psychology (A) and neuroscience (B) by country of affiliation. Figure only include countries represented by ≥1% of editors.

In terms of comparing the 50 journals, at over half (58%) of the top journals in psychology, 50% or more of the editors were affiliated with the USA. Furthermore, at over a quarter of the journals (26%), 75% or more of the editors were affiliated with the USA.

Of the main contributing countries (≥4% of total number of editors in the field), there was a significant difference in the gender distribution of USA-based editors [*X*^2^(1) = 39.7, *p* < .001] which was 58% male and 42% female, UK-based editors [*X*^2^(1) = 22.6, *p* < .001] which was 67% male and 33% female, Canada-based editors [*X*^2^(1) = 10.49, *p* = .001] which was 63% male and 37% female, and Spain-based editors [*X*^2^(1) = 15.9, *p* < .001] which was 68% male and 32% female.

### Neuroscience

#### Gender

The sample included a total of 3,093 editors in the field of neuroscience. Overall, there were significantly more male than female editors (see **Table 2**). This was the case at all editorial levels, with significant differences observed in the proportion of males and females in every category.

**Table 2:** Overall proportion of editors who are male and female in the top 50 journals in the field of neuroscience, and in each of the three sub-categories: 1) Editors-in-chief and their deputies, 2) associate and section editors, and 3) advisory and editorial boards.

| NEUROSCIENCE | Female | Male | Statistic |
| --- | --- | --- | --- |
| Overall (n = 3,085, 8 NA excluded) | 30% | 70% | $\chi^2(1) = 490.3, p < .001$ |
| Editors-in-chief and deputies (n = 171) | 33% | 67% | $\chi^2(1) = 20.4, p < .001$ |
| Associate and section editors (n = 685) | 29% | 71% | $\chi^2(1) = 116.9, p < .001$ |
| Advisory and editorial boards (n = 2,237, 8 NA excluded) | 30% | 70% | $\chi^2(1) = 353.6, p < .001$ |

In contrast to the 88% of neuroscience journals that were comprised of more than 50% male editors, only 10% of journals included a similar proportion of female editors (see **Figure 1B**). 78% of neuroscience journals were comprised of more than 60% male editors, in contrast to 4% of journals that were comprised of a similar proportion of female editors. 40% of journals were comprised of more than 70% male editors, compared with 4% with the same proportion of female editors. 10% of journals had either more than 80% or 90% male editors, compared with 2% and 0%, respectively, that were comprised of the same proportion of female editors. See Supplementary Figure 2 for the binned data by gender at the 50 journals.

#### Geography

As seen in psychology, editors at the top neuroscience journals were primarily based in North America (57%), followed by Europe (29%), Asia (9%), Oceania (4%), Latin America (1%) and Africa (<0.5%). This distribution was significantly skewed towards North America [*X*^2^(5) = 4698.2, *p* < .001]. When we considered the country of affiliation, we found that over half of the editors were based in the United States (52%), followed by the United Kingdom (9%), Germany (7%), and Canada (5%). **Figure 2B** shows the proportion of editors affiliated with each country that contributed more than 1% to the total number of editors analyzed.

In terms of individual differences between journals, at half of the top journals in neuroscience (50%), 50% or more of the editors were affiliated with the USA. At 14% of the journals, 75% or more of the editors were affiliated with the USA. One journal, *Annual Review of Vision Science*, was comprised entirely of USA-based editors.

Of the main contributing countries (≥4% of total number of editors in the field), there was a significant difference in the gender distribution of USA-based editors [*X*^2^(1) = 218.7, *p* < .001] which was 68% male and 32% female, UK-based editors [*X*^2^(1) = 46.7, *p* < .001] which was 70% male and 30% female, Germany-based editors [*X*^2^(1) = 59.7, *p* < .001] which was 77% male and 23% female, and Canada-based editors [*X*^2^(1) = 7.8, *p* = .005] which was 62% male and 38% female.

## Discussion

The present findings reveal that female scholars (as inferred by the authors based on publically available information) are under-represented in editorial board positions in the most popular journals (as indexed by impact factor) in psychology, and even more so in neuroscience. Moreover, in both fields there was a clear and significant geographical over-representation of editors affiliated to the United States in these (English language) journals. The number of editors affiliated with the US in each field was greater than the proportion of journals published in the US, suggesting that even journals published outside the US are skewed in favor of editors from this country. All categories of editor, bar the editors-in-chief of psychology journals, were characterized by significant differences in the proportion of men and women. The situation was widespread, and not driven by a few “bad apples”, with ten times as many journals in neuroscience comprised of more than 70% male editors (40%) compared with the same proportion of female editors (4%). The ratio was similar in psychology, with 22% of journals comprised of more than 70% male editors compared to just 2% that had the same proportion of women. Only at a handful of journals did women visibly outnumber men. Based on these data and the wider literature on academic publishing, one might argue that the ideas, values and decision-making biases of men, particularly those from the United States of America are over-represented in editorial positions of the most recognized academic journals in psychology and neuroscience. We wish to emphasize at the outset, however, that the implications of these findings are limited to inferred gender from publicly available information, not self-reported. Our study also did not address many other traits, identities and motives that may explain and enrich the implications of our data on gender and geographical affiliation, such as for example areas of double disadvantage and intersectionality; the consequences of membership in multiple discriminated against social groups (Cole, 2009).

Of course, academic psychology and neuroscience are fields that have been traditionally dominated by men, and the US has a larger population than the other countries featured in **Figure 2**. It is possible that psychology and neuroscience have also grown to a greater extent as academic disciplines in the US than elsewhere. Moreover, we considered only English-speaking journals, and journals associated with US-based societies were well-represented. While history, affiliation, language and size may explain some of the observed differences, it is of note that the gender ratios of undergraduate and graduate students entering these fields has dramatically changed in recent decades, and most science appears to be shared globally in English. Therefore, to the extent that academic fields wish to avoid restricting the progression and scientific interests of women and non-US based scholars, gender and affiliation diversity of leadership positions in these fields should also change.

Why would the editorial boards of top journals strike to include a heterogenous sample of scholars? Firstly, editorial positions are considered prestigious and influential, and likely impact the career advancement and networking opportunities of those that hold them. Secondly, evidence suggests that equity in science enhances productivity and innovation, and there is evidence that gender equity on editorial boards actually improves the review process (Wing, Benner, Petersen, Newcomb, & Scott, 2010). Thirdly, representation may have a meaningful impact on the next generation of scientists. When undergraduates studying science, engineering and mathematics were randomly assigned to view conference footage that depicted either male-skewed conference attendance (three men to every woman present), or balanced attendance (equal numbers of men and women), the female students who viewed the skewed attendance reported less feelings of belonging than female students who viewed balanced attendance (Murphy, Steele, & Gross, 2007). Male students’ sense of belonging was not impacted by either condition. Fourthly, positive changes in psychology and neuroscience may have the capacity to influence positive change in other academic disciplines.

While it is not possible for the present work to examine the reasons why gender disparity might exist in editorial boards, future qualitative and quantitative studies could shed further light on this matter. For now, there are several possible mechanisms that we wish to highlight. Female tenure-track academics in psychology earn less, publish less, are cited less, and hold fewer grants than their male counterparts (APA Committee on Women in Psychology, 2017; Gruber et al., 2020). These differences may result in women being considered less worthy of positions on editorial boards. Reasons for this reduced productivity may be complex but they seem to include increased childcare demands than male colleagues – American mothers spend on average 75% more time performing childcare duties than fathers (Geiger, Livingston, & Bialik, 2019), and reduced financial resources – male academics in the biomedical sciences receive larger start-up funds than female academics (Sege, Nykiel-Bub, & Selk, 2015).

In general, men and women report similar levels of motivation to engage in mentorship in the workplace (Ragins & Cotton, 1993), however it is unknown whether this extends to journal editing, or reflects the values of men and women working in the fields of neuroscience and psychology specifically. Evidence from the field of political sciences suggests that women faculty were more likely to perform internal service roles (e.g., departmental committee work) while men were more likely to perform higher status external service roles, such as editing (Mitchell & Hesli, 2013). It would be interesting for follow-up work to consider whether female academics are being encouraged and promoted to be editors and/or offered positions on editorial boards at similar rates to their male colleagues, and if so, what factors guide their constrained or unconstrained choices regarding editorial positions (Xie & Shauman, 2003).

### Limitations

In line with similar work in related fields (Amrein et al., 2011; Hafeez et al., 2019; Mariotto et al., 2020), we assigned gender and country of affiliation based on publicly available information, and it is possible that we incorrectly determined some editors’ identities. A more detailed investigation, with institutional ethical approval, would have permitted us to contact editors directly. This would have also allowed us to collect data on the race and/or sexual and identity of editors, and provided a more comprehensive report on the multiple intersecting identifies of editorial teams in psychology and neuroscience. The importance of such work is evidenced by findings of under-representation of academics of color as authors and editors in psychology (Roberts, Baraket-Shavit, Dollins, Goldie, & Mortenson, 2020). Non-heterosexual or gender-conforming academics often feel discouraged from expressing their identity. Of 1,427 science, technology, engineering and mathematics (STEM) academics surveyed who identified as lesbian, gay, bisexual, trans, queer or asexual (LGBTQA) from the United States, United Kingdom, Canada and Australia, fewer than half had disclosed their identity to more than half their colleagues and many had disclosed to few or no colleagues (Yoder & Mattheis, 2016). Thus, it appears that women and non-US based scholars, who also have intersecting racial and/or sexual identities, are likely to experience even less representation on editorial boards.

Finally, the addition of more journals and further methods would have allowed us to capture data in several subfields of psychology and neuroscience, as well as to interview some editors and scientists to understand the current data in greater depth. We nevertheless hope the present work is a useful first step towards such directions. The database generated by *Clarivate Analytics’ Journal Citation Reports* for psychology represents the top generalist psychology journals, and does not capture high impact journals within sub-disciplines of psychology. Future studies should also look at sub-discipline specific journals. Amongst the psychology journals analyzed was *Psychology of Women Quarterly*. Unsurprisingly, the majority of editors at this journal are women. While the presence of this journal in the list is encouraging, it also likely tipped the scales towards greater female editorial representation in psychology.

### Conclusions

Some areas of leadership in neuroscience and psychology are improving in terms of gender parity. The number of female APA presidents, at 70% over the last decade, is the highest it has ever been (Grubber et al., 2020). In other areas, however, women continue to be under-represented, including department chair positions (APA Committee on Women in Psychology, 2017), and, as shown here, representation on the editorial boards of top journals.

It has been suggested that the people who practice science exert significant influence on the types of questions that are asked, the evidence that is collected and analyzed, and the findings that are reported (Leggon, 2006, 2010). We would venture that the over-representation of men, and those affiliated with the USA, in editorial roles at the most influential journals in psychology and neuroscience impacts, and potentially skews, the publication decisions that affect not only the careers of scientists but also the kind of science that gets published in such journals. Future studies should explore the decisions undertaken by editors in this respect. Similarly, whether the over-representation of men and US-based academics noted here disproportionately effects the careers of scientists from under-represented groups remains an open and pertinent question for future research.

In agreement with similar commentaries in related fields (Amrein et al., 2011; Hafeez et al., 2019), we reiterate the call for journals to define their policies and selection criteria for editorial board appointment, and to actively geo-diversify their editorial boards. Some selection criteria, such as publication and citation counts, may themselves be biased against women and those based outside the USA, and should be revised.

## Acknowledgements

This study and AF’s time was supported by the Faculty of Brain Sciences, UCL. We thank Manos Tsakiris and Manjula Patrick for their comments on earlier versions of this paper.

