## Supplementary Materials for "Gender and geographical disparity in editorial boards of journals in psychology and neuroscience"

**Table 1: Categorization of editorial roles.**

| Category 1: Editors-in-chief and their deputies | Category 2: Associate and section editors | Category 3: Editorial board members | Category 4: Advisory board members | Excluded roles |
|---|---|---|---|---|
| Editor-in-chief | Associate editor | Editorial board | Advisory board | Founding editor |
| Co-editor | Reviews editor | Committee member | Editorial advisory board | Editor emeritus |
| Senior editor* | Neurogenesis editor | Editorial committee* | Advisory editor | Student advisor |
| Editor* | Handling editor | Editor* | Honorary advisor | Managing editor |
| Chief editor | Senior associate editor | Junior editor | Consulting editor | Submissions editor |
| Deputy editor | Action editor | Senior editor* | Scientific board | Editorial coadjutant |
| Reviewing editor | Section editor | Editorial review board | Consultant to the editors | Commissioning editor |
| Assistant editor* | Field editor | International editorial board | Supervisory committee | Communications editor |
| Field chief editor | Editorial committee* | | Publication committee | Education editor |
| Senior international editor | International associate editor | | | (Senior) Social media editor |
| | Associate editor for Asia | | | Statistical editor/advisor |
| | Associate editor for North America | | | Senior cultural equity editor |
| | Book review editor | | | Senior expert-by-experience |
| | Teaching section editor | | | Features editor |
| | Editor* | | | Consulting editor for statistics |
| | Assistant editor* | | | Editor of clinical trials highlights |
| | Case editor | | | |
| | Deputy case editor | | | |
| | Specialty chief editor | | | |
| | Senior research editor | | | |
| | Senior consulting editor | | | |

| |
|---|
| Senior psychotherapy editor |
| Associate editor for reviews |
| Scientific editor |

*The same title was used by different journals to denote varying levels of seniority, and so feature in multiple categories. Decisions were made on an individual basis according on the organization of each journal.

**Figure 1: Representation by gender at the top 50 journals in psychology. A) shows the number of journals with the proportion of female (A) and male (B) editors binned by ten-point percentage increments.**

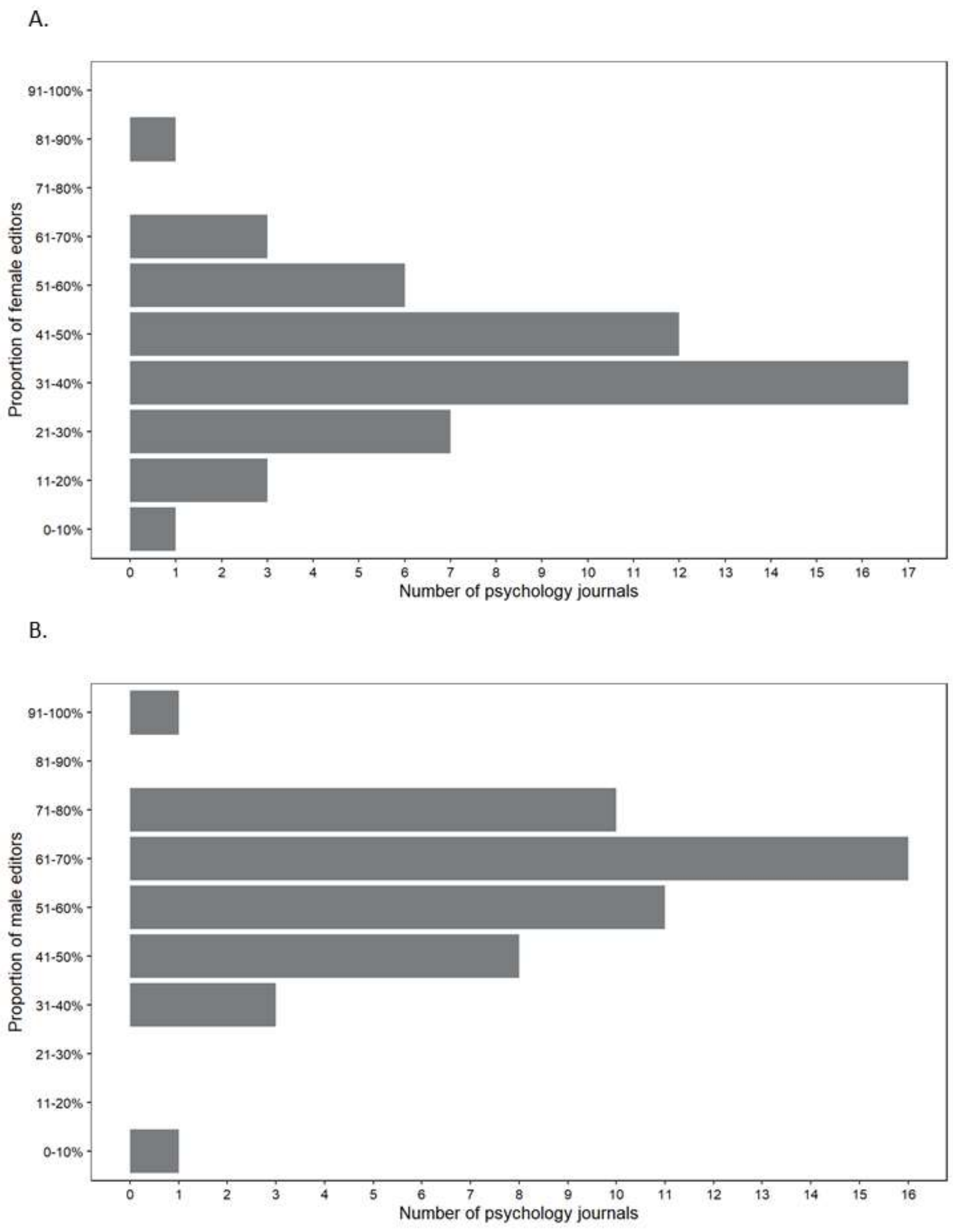

**Figure 2: Representation by gender at the top 50 journals in neuroscience. A) shows the number of journals with the proportion of female (A) and male (B) editors binned by ten-point percentage increments.**

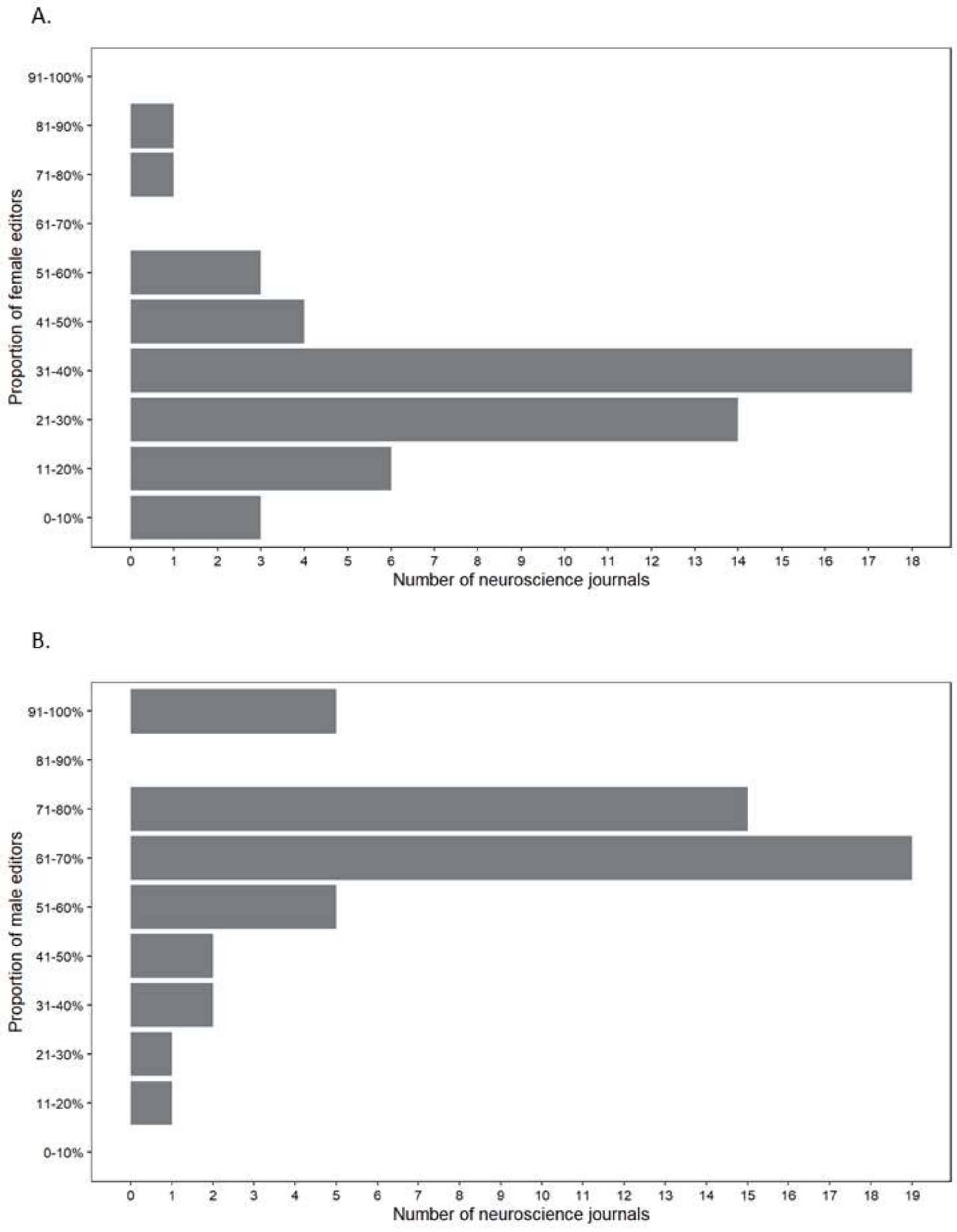

**Table 2: Proportion of editors, in psychology and neuroscience, from all featured countries.**

| Country | Proportion of Psychology editors | Proportion of Neuroscience editors |
|---|---|---|
| United States | 60.58% | 52.43% |
| United Kingdom | 6.89% | 9.18% |
| Spain | 4.99% | 1.03% |
| Canada | 4.64% | 4.53% |
| Netherlands | 3.05% | 1.97% |
| Germany | 3.00% | 6.79% |
| Australia | 2.85% | 3.30% |
| Italy | 1.55% | 1.97% |
| Israel | 1.15% | 0.97% |
| New Zealand | 1.15% | 0.23% |
| Switzerland | 1.00% | 1.62% |
| Austria | 0.70% | 0.55% |
| Hong Kong | 0.65% | 0.32% |
| China | 0.65% | 3.04% |
| France | 0.55% | 2.49% |
| Belgium | 0.55% | 0.68% |
| Ireland | 0.50% | 0.52% |
| South Africa | 0.45% | 0.10% |
| Sweden | 0.40% | 0.81% |
| Denmark | 0.35% | 0.81% |
| Norway | 0.35% | 0.13% |
| Finland | 0.35% | 0.23% |
| Singapore | 0.30% | 0.26% |
| Japan | 0.30% | 2.75% |
| Portugal | 0.25% | 0.16% |
| Poland | 0.25% | 0.10% |
| India | 0.20% | 0.19% |
| Taiwan | 0.20% | 0.23% |
| Greece | 0.20% | 0.13% |
| Mexico | 0.20% | 0.13% |
| Turkey | 0.15% | 0.03% |
| Cyprus | 0.15% | 0.00% |
| Chile | 0.15% | 0.23% |
| Hungary | 0.15% | 0.13% |
| South Korea | 0.15% | 0.10% |
| Estonia | 0.10% | 0.00% |
| Brazil | 0.10% | 0.61% |
| Argentina | 0.10% | 0.19% |
| Lithuania | 0.10% | 0.00% |
| Russia | 0.05% | 0.03% |

| | | |
|---|---|---|
| Bulgaria | 0.05% | 0.00% |
| Ghana | 0.05% | 0.00% |
| Iceland | 0.05% | 0.00% |
| Iran | 0.05% | 0.00% |
| Kenya | 0.05% | 0.00% |
| Nigeria | 0.05% | 0.00% |
| Puerto Rico | 0.05% | 0.00% |
| Algeria | 0.05% | 0.00% |
| Luxembourg | 0.05% | 0.00% |
| Malta | 0.05% | 0.00% |
| Korea | 0.00% | 0.55% |
| Cuba | 0.00% | 0.06% |
| Czech Republic | 0.00% | 0.06% |
| Egypt | 0.00% | 0.03% |
| Lebanon | 0.00% | 0.03% |
| Macao | 0.00% | 0.03% |
| Paraguay | 0.00% | 0.03% |
| Philippines | 0.00% | 0.03% |
| Qatar | 0.00% | 0.03% |
| Romania | 0.00% | 0.03% |
| Saudi Arabia | 0.00% | 0.03% |
| Thailand | 0.00% | 0.03% |
| United Arab Emirates | 0.00% | 0.03% |
| Venezuela | 0.00% | 0.03% |
